# Phage as signatures of healthy microbiomes

**DOI:** 10.1101/2024.03.18.585470

**Authors:** Rachel M. Wheatley, Dominique Holtappels, Britt Koskella

## Abstract

Parasites are foundational to ecosystem health both as indicator species of community productivity but also as drivers of diversity. In bacterial communities, bacteriophage viruses can play such a role as they track the dynamic composition of bacterial hosts, and in the case of lytic phages, confer a growth advantage to lower abundance bacteria while adapting to more common ones. We set out to test whether viromes can be used as signatures of microbiome health using previously published results across systems. By comparing observed phage and bacterial diversity between microbiomes characterized by disturbance (so-called dysbiosis) and those considered control populations, we were able to identify some key commonalities. While just under half of studies report significant changes in viral species richness in dysbiosis, just under two thirds of studies find the viral composition to shift in dysbiosis, with specific viral taxa enrichment acting as a common signature of dysbiosis. Our analyses also suggest that the positive relationship between bacteriome and virome alpha diversity observed in health breaks down under microbiome disturbance. Overall, while specific viral signatures of dysbiosis are likely to be highly disease- and condition-specific, existing ecological theory shows clear promise in predicting and explaining microbiome health. Future data on bacteria-phage diversity relationships may provide us with much needed opportunity to diagnose, treat, and better understand the causes of dysbiosis.

**Research in context:** *Evidence before this study:* Being able to identify signatures of microbiome health (or lack thereof) has the potential to improve the way we diagnose and treat disease. To do this, the bacterial microbiome is traditionally characterised at the 16S taxonomic level, and changes in composition are linked to changes in disease status. More recently, the field of viromics has gained attention, and studies have begun to probe the relationship between the virome and health or disturbance (‘dysbiosis’). This work has focused to date on finding single phages that indicate presence of known pathogens, or in a few cases the relationship between viral diversity and disease. To our knowledge, no work has yet sought to identify a common signature of dysbiosis or find commonalities across systems that suggest a role for phages in dysbiosis. Decades of ecological theory has shown how parasites can shape the ecology and evolution of their hosts, and here we argue that bacteriophage viruses have the potential to shape these same processes within microbial communities. The motivation for the current work was thus to ask whether existing ecological theory could help us identify viral signatures of dysbiosis in the microbiome.

*Added value of this study:* This study employed a systematic review and meta-analysis to test whether and when phage communities can be used as signatures of microbiome health. To do this, we synthesized previously published results that measure composition of the virome between bacterial microbiomes characterised by health or dysbiosis. We found a total of 39 studies across human, mouse, pig and cow hosts that spanned a diverse spectrum of dysbioses, including bacterial infections, viral infections, and varied diseases such as cancer, cirrhosis, and inflammatory bowel disease, and identified a number of commonalities. Just under half of these studies reported a significant change in viral species richness in dysbiosis, and just under two thirds reported the viral composition to shift in dysbiosis. While the vast majority of studies report an enrichment of specific viral taxa associated with dysbiosis, there is little overlap among studies regarding the identity of these enriched taxa. Finally, our analysis provides evidence that the positive relationship between bacteriome and virome alpha diversity breaks down in dysbiosis.

*Implications of all available evidence:* Synthesis of the available evidence suggests that while looking for specific viral taxa as signatures may be limited to associations that are highly disease or condition specific, there is promise for the use of existing ecological theory in predicting and explaining microbiome health when considering compositional changes in the virome. Prospective studies should look to expand the data we have on bacteria-phage relationships at the level of species richness and community compositions, and we argue that more routinely investigating the virome or phageome, in addition to collecting 16S taxonomic descriptions of the microbial community, would help improve our ability to identify signatures of microbiome health. These viral signatures may offer early warning signs of microbiome disturbance and disease. This has clear relevance to our ability to diagnose, treat, and understand the underlying causes of disease.

## Introduction

Linking species diversity to community productivity and ecosystem function is a central tenet of Ecological research. These ideas have also been recently co-opted to help explain, and ultimately manipulate, the diversity of host-associated microbiomes (*1, 2*). By considering coexistence of microbial species as a result of interactions among them (e.g. competition, cross-feeding, or parasitism), ecological theory has offered a highly useful guide to explaining microbiome diversity (*3, 4*). As our sequencing and ‘omics capabilities continue to expand, the utility and necessity of these models has been further clarified (*4*). In particular, viral metagenomic approaches have offered new opportunities to explore the role of parasites in shaping microbiome stability, diversity, and function (*5*).

In their insightful review from 2006, Hudson and coauthors posed the question: “Is a healthy ecosystem one that is rich in parasites?” (*6*). They posed this based on the idea that parasites and pathogens shape the ecology and evolution of their hosts, including through modulating competition and energy flow, and thus can act as key drivers of biodiversity (*7-10*). When there are thriving and diverse host populations, we expect thriving and diverse parasitic ones. Higher population densities of hosts should result in higher parasite abundances (*11*); a relationship that has been well-documented in plant and animal communities (*12-15*). Given the inherent reliance of parasites on their hosts, and the threshold host density under which parasite populations generally go extinct before host populations (*16*), parasite diversity loss could be more pronounced than host loss and thus act as an early warning sign of disturbance/disfunction, as supported by theoretical models (*17, 18*).

In the microbial world, bacteriophage viruses (phage) have the potential to act as signatures and as drivers of bacterial community diversity (*19*). We therefore ask: is a healthy microbiome one that is rich in phages? To answer this question, one first needs to differentiate a healthy microbiome from a disturbed one (often referred to as dysbiosis). Hudson and colleagues followed the logic of Costanza and Mageau (1999) to define a healthy ecosystem as “one that persists, maintains vigor (productivity), organization (biodiversity and predictability) and resilience (time to recovery)” (*6, 20*). Similar ‘rules’ have been applied to describing a healthy microbiome (*21, 22*). However, it is more straight-forward to focus on microbial communities that are associated with a diseased state, where changes in microbiome composition, diversity, and/or function can be linked to disease (either via correlational change or causation). These dysbioses are ideal situations in which to test whether phage prevalence, composition or diversity relates to microbiome health (or lack thereof).

If we define microbiome dysbiosis as a compositional alteration in individuals with disease compared with healthy subjects, we can then consider the following as possible mechanisms: (1) an overgrowth of key taxa (e.g. excessive growth of potentially harmful microbes), (2) a change in diversity (often understood as a loss of diversity, for example in the human gut (*23, 24*)), (3) a loss of key taxa (e.g. loss of potentially beneficial microbes), and/or (4) altered microbiome function (*25-27*). In each case, viromes may provide useful signatures of these dynamics (Figure 1). For example, a change in bacterial diversity may in turn result in a change in phage diversity (*28, 29*), an overgrowth of bacterial taxa may result in an enrichment of specific phage taxa (*30*), and a loss of bacterial taxa may result in a depletion of specific phage taxa for which the bacteria are hosts. Microbiome dysbiosis in many disease states have been associated with an overall loss of bacterial diversity (*23, 24, 31*), and as such, one might predict a loss of virome diversity to also be a common signature in dysbiosis (*28*).

**Figure 1.**
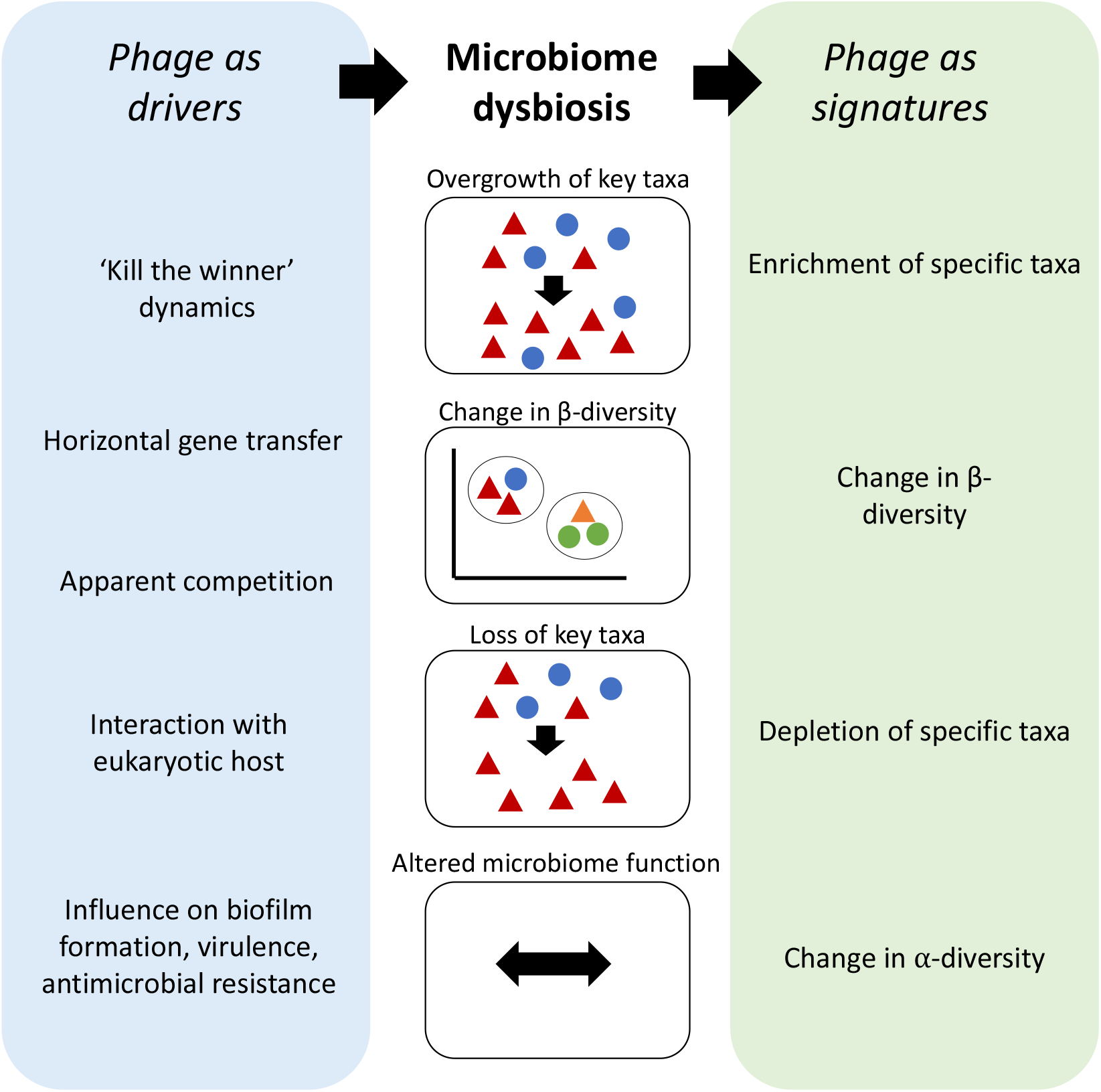
Framework outlining the putative relationship between microbiome dysbiosis and phage as signatures or drivers of these shifts.

Beyond acting as signatures of microbiome health, phage may also act as drivers of dysbiosis (Figure 1). For example, both lytic phages (those requiring cell lysis for onward transmission) and temperate phages (those that can stably transmit among bacterial cells during cell division) can alter competitive outcomes among bacterial strains both via direct lysing of cells and by selecting for costly phage resistance mechanisms (*19, 32*). Lytic phages can reduce population sizes of highly abundant and competitive bacterial species, opening up key resources for more rare bacterial types via ‘kill the winner’ dynamics (*33*). Temperate phages can drive changes in microbiome function and composition through horizontal gene transfer (HGT)(*19*), providing new functions to bacterial hosts that modulate bacteria-bacteria competition and influence how bacterial hosts respond to exogenous selective pressures such as antibiotics (*34*). Finally, recent evidence has highlighted that phages can directly modulate mammalian host immunity, in turn driving changes in microbiome composition (*35*).

Although the idea that phages could play a role in gut health has been around for decades, with early evidence from coliphages suggesting increased lytic phage activity in intestinal diseases (*36*), the true utility of this approach has been beyond reach due to both the costs of generating and difficulty of analyzing viromic datasets. As our understanding of microbiome dysbiosis across disease systems continues to expand, there is new opportunity to move beyond pairwise bacteria-phage interactions to take a more holistic approach to understanding the role of phages in microbiome health. Here, we systematically assessed studies quantifying viral diversity across healthy and disturbed systems to test whether phage diversity, the relationship between phage and bacterial diversity, and/or specific phage compositions were typically altered across these states. We sought to determine whether viromic data can be used as a marker of current or pending microbiome disturbance, as well as asking how such signatures might depend upon the system, type of dysbiosis, and profiling or data analysis method.

## Methods

To identify studies in which the virome was compared between a healthy state and a dysbiosis state and a measure of virome diversity (alpha or beta) was obtained, we focused on studies that compared samples associated with disease, infections, interventions, or perturbations to a control group with a defined absence of such dysbiosis. We used a combination of keyword and backwards and forwards citation searches in both PubMed and Web of Science using a combination of search terms; ‘virome’, ‘virus’, ‘phageome’, ‘phage’, ‘disease’, ‘diversity’, ‘dysbiosis’, ‘infection’ with backwards and forwards citation searches conducted in parallel. The final search was conducted in December 2022. A total of 39 studies were identified that met the criteria for inclusion (Supplementary Methods)(Supplemental Table 1), each of which was characterized based on: (1) system (e.g. host organism and microbiome site), (2) type of dysbiosis, (3) method of viral isolation, (4) virus sequence analysis pipeline, (3) host age, (4) virome alpha diversity, (5) virome beta diversity, (6) changes to viral taxa (e.g. enrichment of specific OTU), and (7) classification of viral diversity (e.g. on phage or on whole virome) (Supplementary Table 1, Supplementary Table 2). Initially, we collected and analysed all dysbiosis comparisons within a study that corresponded to independent samples (e.g. either different dysbiosis groupings, different timepoints, different microbiome sites, different treatments, or different methods of viral isolation; Supplementary Table 1). We then filtered this table to one comparison per study to limit possible pseudoreplication (Supplementary Table 2), filtering to select for the comparison that reported the largest difference in virome diversity (regardless of direction). We took two approaches to analyse this qualitative data: (1) an analysis of one comparison per study (n=39 studies), and (2) an analysis of all separate comparisons per study (n=70 comparisons from 39 studies). JMP Pro (v.16.0.0) was used to build contingency tables to evaluate the effect between the different variables with Fisher’s exact test (*37*).

The majority of the 39 studies that met our criterion for inclusion focused on human hosts (30 studies (*29, 38-66*)), with the remaining studies using cow (1 study (*67*)), pig (1 study (*68*)), or mouse (7 studies (*69-75*)) (Supplementary Table 2). These studies covered a diverse spectrum of dysbioses, spanning cancer to viral infections such as HIV and COVID-19, and varied conditions such as alcohol hepatitis, periodontal disease, spinal injury, bacterial vaginosis, cirrhosis, obesity, rheumatoid arthritis, inflammatory bowel disease. The majority of studies investigated the virome in adults (n=32) and used viral isolation that targets free viral particles present in a sample (e.g. through filtration and concentration steps; n=30). The remaining studies (n=9) used whole sample metagenomics, meaning the viruses associated with the bacterial fraction (e.g. prophages) will also be represented in the sample. Some studies filtered their samples and data to focus specifically on the phageome (n=15), whereas others describe the whole virome (n=24).

From the full dataset, we were able to obtain virome alpha diversity data (shannons diversity scores for all individuals within the study) in dysbiosis and health from a subset of 16 studies. This was done either through supplementary/publicly available data files, correspondence with authors, or graph extraction. For graph extraction, we used the web-based tool WebPlotDigitizer to extract numerical data from plot images (*76*). For studies with multiple dysbiosis group comparisons, all data was initially collected where available, but we again sought to limit the impact of pseudoreplication in the analysis by condensing to one data comparison per study where possible (Supplementary Table 3). For example, when multiple timepoints were assessed within a study, the timepoint closest to dysbiosis was chosen and when multiple dysbiosis severities were assessed, the most severe was chosen. This resulted in a total of 20 data comparisons on virome alpha diversity in control compared to a dysbiosis, from across 16 studies with which we performed a meta-analysis. The meta-analysis was conducted following Borenstein et al. (2009)(*77*), using a similar approach as used in (*78, 79*), and is described in (Supplementary Methods)(Supplementary table 3).

A subset of the studies (n=24) also analyzed the bacteriome within the same samples as the virome via 16S sequencing or metagenomes. For three of these studies, the paired bacteriome analysis was presented in a separate paper (*80-82*). For our qualitative analysis, we recorded whether the study reported a significant decrease (-1), no change (0) or significant increase (+1) in alpha diversity between healthy individuals and those characterized by dysbiosis (Supplementary Table 1) and for the beta diversity we recorded whether the study reported either no shift in beta diversity (0) or a significant shift in diversity (+1) (Supplementary Table 2). We analyzed the relation of the different parameters by means on contingency tables using the JMP Pro (*37*). From the 16 studies included in the meta-analysis of virome alpha diversity, we were able to obtain bacterial alpha diversity datasets paired with the virome data from a total of 7 studies and 9 data comparisons (Supplementary Table 4). To determine the correlation between paired bacterial alpha diversity and viral alpha diversity, and to what extent bacterial diversity was predictive for viral diversity, we calculated Pearson correlation coefficient (*r*) and the coefficient of determination (*r*^2^) for these paired datasets using R (Supplementary Table 4) (*83*).

## Results

The majority of studies in our dataset (n=35/39) investigated changes to alpha diversity of the virome in dysbiosis (Supplementary Table 2), out of which 8 studies (23%, n=8/35) report alpha diversity to decrease in dysbiosis, 8 studies report alpha diversity to increase in dysbiosis (23%, n=8/35), and the remaining 19 studies (54%, n=19/35) reported no significant difference to alpha diversity in dysbiosis. For the mouse and human gut, we found a significant difference in the relationship between alpha diversity and dybiosis (Chisq p-value 0.0138) (Supplementary Table 2). In contrast to the virome alpha diversity of the mouse gut which increased in dysbiosis (n=4/7 – 67%), the virome alpha diversity in the human gut did not change, with only 9% of manuscripts reporting an increase of the alpha diversity (n=2/22 studies). For the 20 datasets in which we obtained individual virome alpha diversity values, just over half of the datasets (11/20) gave a negative response ratio for alpha diversity and just under half (9/20) gave a positive response ratio for alpha diversity, with the majority of 95% confidence intervals crossing zero (where zero corresponds to no change in alpha diversity (Figure 2). The summary response ratio was overall determined to be different from zero and very weakly, yet significantly, positive (summary response ratio: 0.031 ± 0.012, two tailed t-test p=0.010) (Supplementary Table 3; Figure 2). These results were not affected by the metagenome used (i.e. VLP versus whole metagenome; one-tailed unpaired t-test; p=0.497), nor between dysbiosis caused by a pathogen versus those where no pathogen was implicated (one-tailed unpaired t-test; p=0.244).

**Figure 2.**
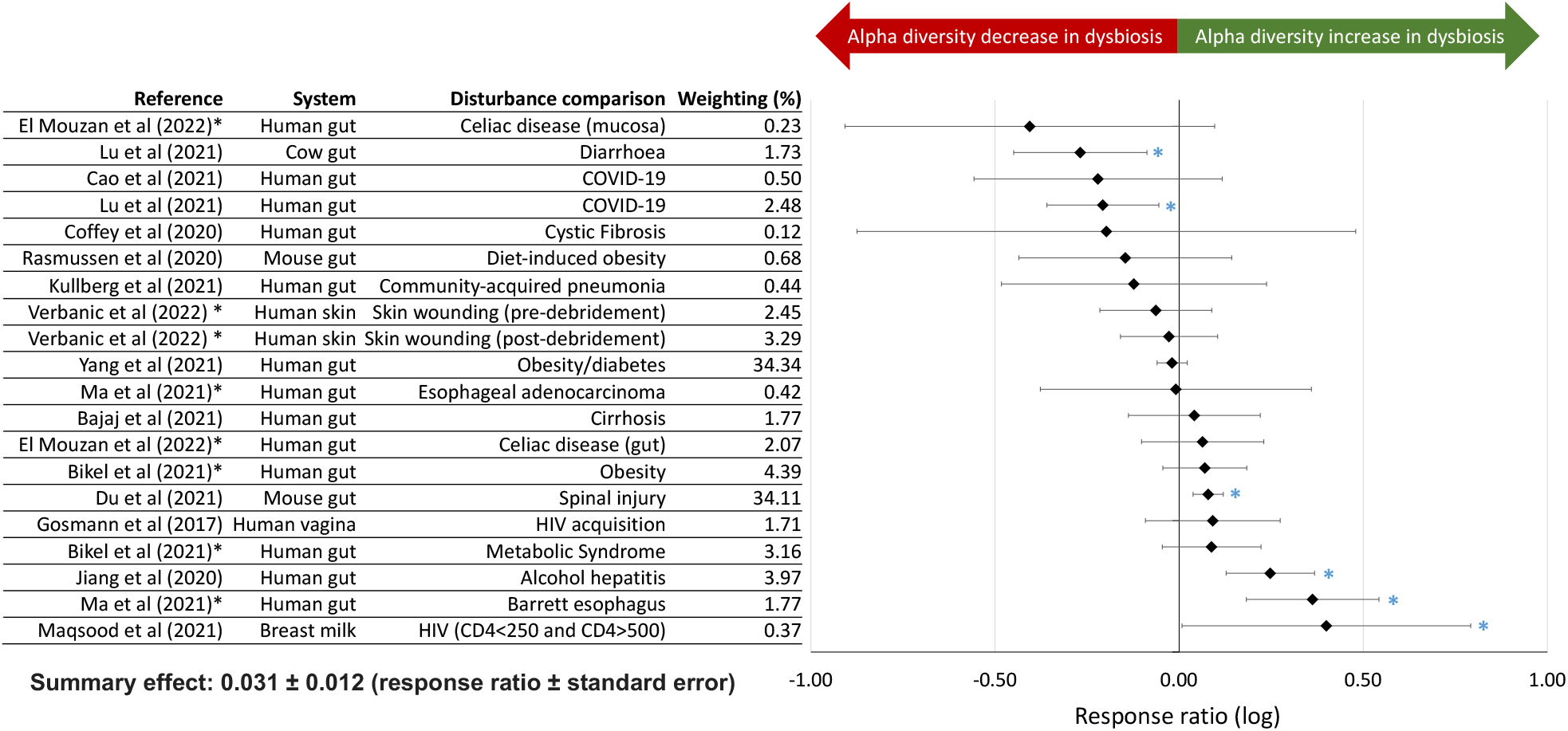
Changes in virome alpha diversity measured as response ratio (log). Positive or negative values indicate where virome alpha diversity increases or decreases in dysbiosis, respectively. A response ratio of zero indicates no change in virome alpha diversity. The bars show the 95% confidence intervals, and blue asterisks (*) indicate where effects have 95% confidence intervals that do not cross zero. References in the table marked with an asterisk (*) indicate studies which have contributed two datasets to the analysis. Weighting of studies for the summary effect are indicated in the table.

Out of studies characterizing beta diversity (composition) of the virome (n=35/39), the majority (63%) report a shift in beta diversity in dysbiosis (Supplementary Table 2). While we again found no significant effects of isolation method (targeted or whole sample metagenomics) or classification of viral diversity used (phageome or virome analysis), we did find a significant difference among studies on the mouse versus human gut (Fisher’s exact test p-value 0.0227). While all mouse gut studies described a shift in beta diversity, only 47% of human gut studies described a similar shift (Supplementary Table 2). Of the studies investigating both alpha and beta diversity, the majority (n = 18, 58%) reported no change in alpha diversity but even in these cases 50% reported a shift in beta diversity. Where richness of viral communities decreases in dysbiosis, a shift in the community composition was always observed and when community richness increased in dysbiosis, 71% of cases also observed a shift in beta diversity.

Of studies testing for enrichment of viral taxa (n=36/39), the majority (83%) reported a significant enrichment of specific viral classes under dysbiosis (Supplementary Table 2). This included 85% of studies using targeted metaviromics and 78% of studies using whole sample metagenomics. Similarly, 87% of phageome studies and 81% of virome studies observed enrichment. Even when the overall richness of the viral community is unchanged or even decreasing in dysbiosis, 82% and 87.5% of the manuscripts still reported an enrichment effect of certain taxa, respectively. When we include all sample comparisons within a study (n=70 comparisons from 39 studies;

Supplementary Table 1), out of the comparisons that investigated viral taxa enrichment (n=60), the majority (83%, n=50/60) report an enrichment of specific viral taxa in dysbiosis.

While enrichment at a fine scale was not comparable across studies due to differences in viral discovery pipelines used, we were able to examine phage enrichment across studies at the family level. Of the nine studies in which crass-like phages were found to differ with disturbance, eight were found to have lower abundance in dysbiosis. Studies with significant enrichment of *Microviridae* were evenly split as enriched in health (5) and disturbance (5), and this was similar for studies finding enrichment of *Siphoviridae* (12 lower and 14 higher in disturbance). Of the 19 studies finding enrichment of *Myoviridae*, 63% described enrichment in dysbiosis (compared to 37% describing enrichment in healthy populations). Of the nine studies identifying enrichment of *Podoviridae*, only two described lower abundances in disturbance (with 7 describing enrichment in disturbance).

Twenty-three studies in our dataset reported both bacterial and viral alpha diversity (Supplementary Table 2), with half reporting a decrease in bacterial alpha diversity with dysbiosis (n=12/23), 17% reporting an increase in dysbiosis (n=4/23), and 30% finding no significant change (n=7/23). Of the 12 studies reporting decreased bacterial alpha diversity in dysbiosis (n=12/23) eight report no correlated changes in viral alpha diversity. Sixteen studies reported both bacterial and viral beta diversity, and the majority of these report a shift in bacterial beta diversity in dysbiosis (75%). In these cases, viral beta diversity typically also shifted in dysbiosis (83% of studies).

Finally, for the nine datasets in which we could acquire quantitative paired data on alpha diversity for both the virome and bacteriome, we found Pearson correlation coefficient (*r*) to be highly variable in both dysbiosis and health (Supplementary Table 4). Control groups showed a weakly positive correlation (mean *r* = 0.2214) which was higher than that of dysbiosis groups (*r* = 0.0400), but this difference was not statistically significant (one-tailed paired t-test, p=0.168). Parallel calculations of the coefficient of determination (*r*^2^) suggested that bacterial alpha diversity is a greater predictor of virome alpha diversity in healthy groups (mean *r*^2^ = 0.3109) than dysbiosis groups (mean *r*^2^ = 0.0839; one-tailed paired t-test, p=0.046) (Figure 3). For the three (out of 9) datasets in which *r*^2^ was higher in dysbiosis than in health, the disturbance examined was non-pathogen related (n=3/3) and nutrition-related (n=3/3).

**Figure 3.**
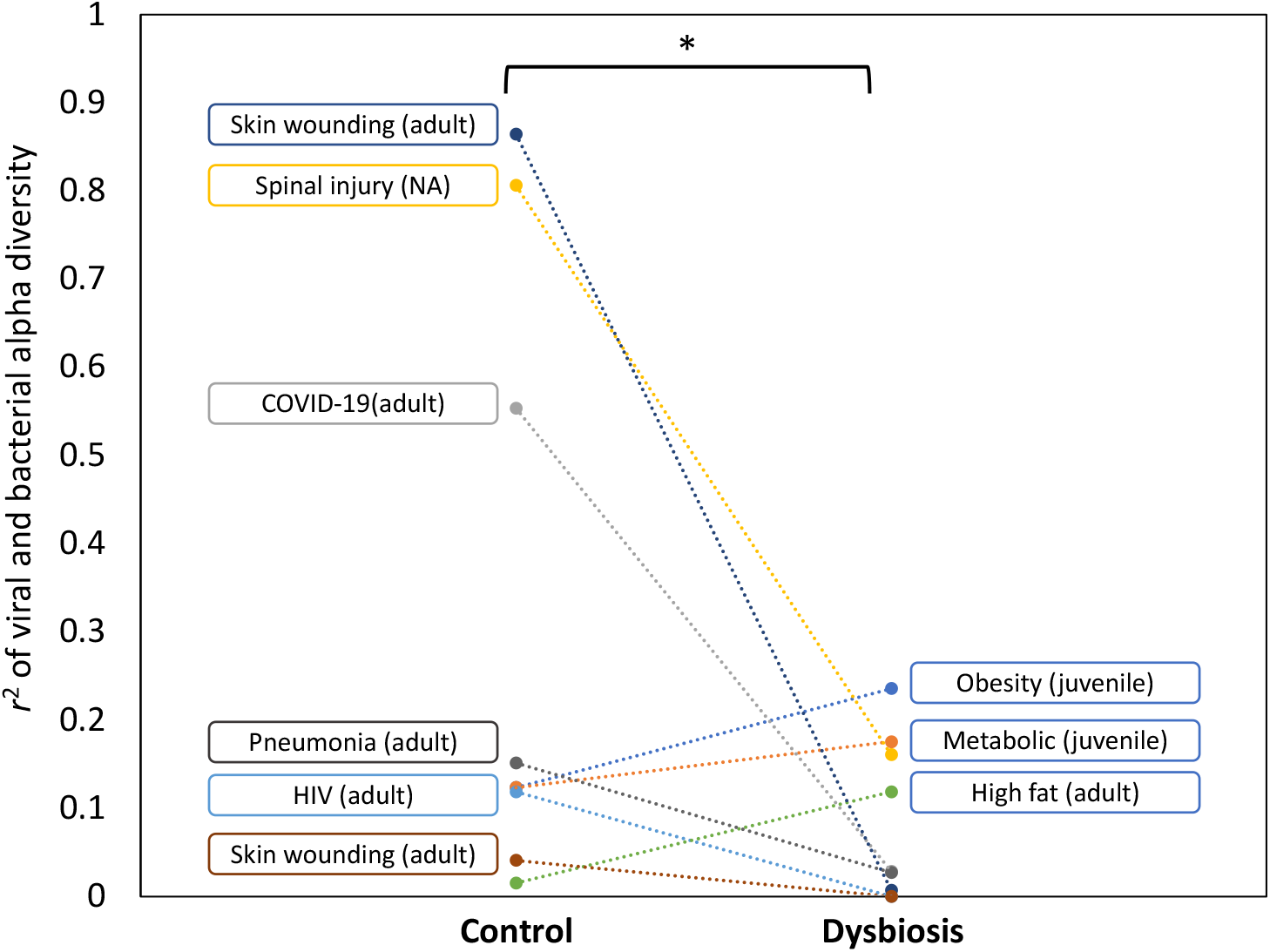
The coefficient of determination (r^2^) between alpha diversity of the viral and bacterial samples from the 9 datasets extracted from 7 studies. Studies are labelled with dysbiosis type and host age in the comparison (control or dysbiosis) in which r^2^ was larger. This suggests bacterial alpha diversity is a greater predictor of viral alpha diversity in control rather than dysbiosis conditions (paired one-tailed t-test, p=0.046). NA; non-available (host age).

## Discussion

There is clear need to uncover signatures of microbiome dysbiosis that move beyond 16S taxonomic descriptions, especially those that can be used either as early detection or applied across a wide range of conditions. We borrow from classic ecological theory of how parasites reflect and impact ecosystem function to ask whether phages act as signatures of microbiome health (*6-8, 11*). The explosion of viromics research in recent years has opened new possibilities for using phage sequences as indicators of pathogen presence (e.g. (*30*), but relatively few studies have yet sought to quantify virome diversity across health and disease. To leverage this first cohort of studies with an aim towards guiding future efforts, we take a conservative approach using author-reported diversity metrics across a broad spectrum of systems and disturbances to search for general signatures of dysbiosis. We recognize several issues that could affect comparison of viromics data across systems, including host developmental stage, collection and filtration methods, data analysis pipelines, and type of disturbance examined (e.g. pathogen-mediated versus non-pathogen related), and therefore included these factors in our analyses. However, analyses such as this are inherently limited by natural variation among individuals (*84-86*), the relative compositions of virulent, lytic phage versus those with a temperate lifestyle, and existing limitations of viral taxonomic databases. Despite these caveats, we observe some striking results that suggest phages hold great promise as signatures of dysbiosis across systems.

First, we find that only 16 out of 25 studies in our dataset report significant change to alpha diversity in dysbiosis (Supplementary Table 2), and that the directional change of alpha diversity is remarkably variable (Figure 2, Supplementary Table 3) despite over half of studies reporting a decrease in bacterial alpha diversity with dysbiosis. In contrast, we find shifting virome composition to be a more consistent signature of dysbiosis, with 63% of studies reporting a significant change in beta diversity with dysbiosis, similar to that observed for the bacterial microbiome. Moreover, the vast majority of studies (83%) report an enrichment of specific viral taxa under dysbiosis. For example, 8 out of 9 studies observed crass-like phages to be less abundant in dysbiosis, in line with a recent in-depth multi-omics study of the viromes associated with IBD (*87*). These data highlight the budding use of specific phage markers for disease, but also indicate that viral community data might also be useful as general markers of disturbance.

Our second key finding is that the relationship between bacteriome and virome alpha diversity breaks down in dysbiosis compared to health, as the bacteriome loses predictive power for the virome after disturbance (Figure 3). This was especially clear in pathogen-associated dysbioses, and we in fact observed an increased correlation in dysbiosis than in health in cases where the disturbance examined was non-pathogen / nutrition-related. A positive correlation between alpha diversity of the bacteriome and virome has previously been demonstrated in the healthy human gut (*28, 29*). That such a relationship may weaken in dysbiosis could help explain the variability we see in directional change of virome alpha diversity. This has implications for our ability to use virome alpha diversity metrics in the same way as bacteriome alpha diversity metrics have emerged as a common signature of human gut dysbiosis (*23, 24*), but leaves open the possibility that shifting viral diversity could be an early warning sign of, or even cause of, bacterial dysbiosis. For example in the case of *Pseudomonas aeruginosa*, the presence of phage Pf has been demonstrated to increase the pathogenicity of the host, its competitive interaction in a microbial community, biofilm formation, and antibiotic resistance, as well as reduced lung function in patients with Cystic Fibrosis (*88-90*). Similarly, the presence of phage phiCTX in *Vibrio cholerae* alters the bacterium’s virulence drastically (*91*).

In sum, we suggest that existing ecological theory on the role parasites play in ecosystems, including as signatures of disturbance and drivers of diversity, is likely to be highly useful in predicting or explaining microbiome health and function. Considering the currently limited data and variation in methods being used, however, there remain a number of open questions. First, in light of the threshold host density under which parasite extinction will occur before its host population (*16*), it is worth exploring whether these viromic signatures act as early warning signs of dysbiosis. Although we currently lack the high-resolution temporal data to dissect this across systems in full, future efforts focused on this question could be highly fruitful. Moreover, as we expand both the number of datasets and types of dybioses being examined, it will be increasingly possible to determine whether course scale signatures that we begin to see here solidify further. For example, in terms of the breakdown in relationship among bacterial and viral diversity, or the seeming shift from temperate phage to lytic ones after disturbance. Such signatures would allow for detection of, and eventually correction of, disturbances in the absence of specific markers of disease or known syndromes. Finally, beyond acting as signatures of dysbiosis, there is clear and emerging need to better understand the direct role phage communities play in shaping microbiome health in order to leverage their promise as therapeutics.

## Supplementary Information

Supplementary Table 1. Dataset of 39 studies that met the inclusion criteria, listing all comparisons from discrete samples.

Supplementary Table 2. Dataset of 39 studies filtered to one comparison per study.

Supplementary Table 3. Meta-analysis of virome alpha diversity.

Supplementary Table 4. Analysis of alpha diversity in the bacteriome and virome.

## Acknowledgements

We would like to thank all authors who publicly shared their datasets in order to advance scientific progress, but in addition would like to thank the following individuals for going above and beyond and sharing further data or information as requested: Prof Adrian Ochoa-Leyva, Prof Jun Wang, Prof Lori Holtz, Dr Robert Kullberg, Dr Xiaolong Liang, Prof Mark Radosevich, Prof Ruth Ley, Dr Leonardo Moreno, Prof Phillip Popovich, Prof Matthew Sullivan, and Dr Ahmed Zayed, Prof Scott A. Handley, Prof Douglas S. Kwon, Dr Torben Rasmussen, Dr Dennis Nielsen, Dr Mohammadali Khan Mirzaei, Dr Corinne Maurice, Dr Colin Hill, and Dr Adina Howe. R.M.W was supported in this work through the George Grosvenor Freeman Fellowship by Examination in Science, Magdalen College (Oxford). R.M.W and B.K. were further supported by fellowships at the Wissenshaftskolleg zu Berlin. D.H. is supported by NSF CAREER award (NSF # 1942881, USDA/NIFA # 1024053). B.K. is a Chan Zuckerburg San Francisco BioHub investigator.

